# αGalCer/CD1d treatment induces pro-inflammatory iNKT1-associated immune responses without aggravating doxorubicin-induced cardiotoxicity

**DOI:** 10.1101/2025.09.18.676932

**Authors:** Sarojini Singh, Roopa Hebbandi Nanjundappa, Harish Babu Kolla, Deepak S. Chauhan, Aahana Palyada, Prasanna Krishnamurthy, Channakeshava Sokke Umeshappa

**Affiliations:** Department of Biomedical Engineering, Heersink School of Medicine and School of Engineering, University of Alabama at Birmingham, Birmingham, AL, United States; Department of Microbiology and Immunology, Dalhousie University, Halifax, Nova Scotia, Canada

**Keywords:** Invariant natural killer T (iNKT) cells, iNKT subsets, Doxorubicin-induced cardiotoxicity, Myocarditis, Alpha Galactosyl ceramide/cluster of differentiation 1d (αGalCer/CD1d), Interferon-γ (IFN-γ)

## Abstract

Invariant natural killer T (iNKT) cells are a unique subset of innate immune cells activated by T cell receptor (TCR) signaling through CD1d molecules presenting lipid antigens. By localizing to specific tissues, iNKT cells play critical roles in maintaining homeostasis and regulating pathophysiology, making them attractive therapeutic targets for inflammatory diseases. However, their frequency in the heart and potential roles in mitigating inflammation-induced cardiac damage remain poorly understood. In this study, we examined the distribution of cardiac iNKT subsets and their response to TCR-directed alpha Galactosyl ceramide/cluster of differentiation 1d (αGalCer/CD1d) complexes in a mouse model of Doxorubicin (Dox)-induced cardiotoxicity. In healthy mice, the heart harbored distinct iNKT subsets, dominated by pro-inflammatory iNKT1 cells, followed by iNKT2 subsets. Treatment with αGalCer/CD1d complexes preferentially expanded the iNKT1 subset. In Dox-induced cardiotoxicity, this treatment unexpectedly increased both IFN-γ, a hallmark cytokine of iNKT1 cells, and IL-10, a key anti-inflammatory mediator typically associated with regulatory iNKT subsets. Despite the expansion of iNKT1 cells and elevated IFN-γ levels, αGalCer/CD1d treatment did not exacerbate cardiac damage and showed modest clinical improvements compared to controls. These findings highlight the complexity of iNKT-mediated immune regulation in the heart and emphasize the need for modified TCR-directed immunotherapeutics to skew immune responses toward a regulatory phenotype. Such approaches could hold promise for treating inflammatory cardiac diseases, including Dox-induced myocarditis, myocardial infarction, and autoimmune myocarditis.

## 1 Introduction

Invariant natural killer T (iNKT) cells occupy a unique niche at the interface of the innate and adaptive immune systems, displaying features characteristic of both natural killer (NK) cells and T cells (1, 2) These cells express a semi-invariant T cell receptor (TCR) composed of an invariant TCRα chain (Vα14-Jα18 in mice; Vα24-Jα18 in humans) paired with a limited repertoire of TCRβ chains (Vβ8, 7, or 2 in mice; Vβ11 in humans) (3–6). iNKT cells recognize lipid antigens presented by the non-polymorphic major histocompatibility complex (MHC) class I-like molecule CD1d, enabling rapid cytokine and chemokine secretion that profoundly shapes immune responses (6–10). Through their tissue-specific localization, iNKT cells contribute to the homeostasis and pathophysiology of various organs, making them promising targets for immunotherapy in inflammatory diseases.

iNKT cells are categorized into several subsets based on their transcription factor expression and cytokine secretion profiles, paralleling conventional CD4+ T cell subsets. These include iNKT1 (PLZFlow, T-bet+, IFN-γ-secreting), iNKT2 (PLZFhi, IL-4-secreting), iNKT17 (PLZFint, RORγt+, IL-17-secreting), iNKTR/iNKT10 (E4BP4+ or Foxp3+, IL-10-secreting), and follicular helper iNKT (iNKTFH, BCL-6+, IL-21-secreting) (11, 12). Each subset exhibits tissue-specificity and preferential homing (13). For instance, iNKT1 predominates in the liver and spleen, iNKT2 in the lungs and intestine, and iNKT17 in the skin, while iNKT^FH^ localizes to lymph nodes and lungs (11, 12). Furthermore, regulatory iNKT subsets—such as adipose-specific iNKT10 (14–16), Breg- induced iNKT cells in joints (17), iNKTR in cutaneous and nervous tissues (18, 19), c-Maf+ iNKTR1 in liver tissues (6, 20), and Foxp3+ iNKTR cells in lymph nodes (21, 22)—suppress inflammation driven by conventional T cells and antigen-presenting cells (APCs). Despite this diversity, the predominant iNKT subsets in the heart and their potential roles in cardiac health and disease remain poorly understood. Whether these subsets can be therapeutically manipulated in cardiac inflammatory diseases remains an open question.

Doxorubicin (Dox), a potent antineoplastic agent, is widely used to treat hematological malignancies and solid tumors. However, its clinical utility is significantly limited by dose-dependent cardiotoxicity, characterized by cardiomyocyte death and progressive cardiac dysfunction (5, 23, 24). Dox-induced cardiomyopathy is mediated in part by the release of pro-inflammatory cytokines, including IL-1β, IL-6, and TNF-α, which drive inflammation and myocardial injury (25, 26). Current strategies to mitigate these complications are largely palliative, underscoring the urgent need for effective therapies. Intriguingly, reducing pro-inflammatory cytokines while enhancing anti-inflammatory mediators, such as IL-10, has shown promise in experimental models of Dox-induced cardiomyopathy (25, 26).

Therapeutic interventions targeting tissue-specific autoimmune diseases with soluble peptide-MHC complexes have demonstrated efficacy through mechanisms such as autoreactive clonal deletion, expansion of antigen-specific regulatory T cells, and induction of IL-10 production (27–29). Similarly, the administration of soluble αGalCer-loaded CD1d complexes (αGalCer/CD1d) has emerged as a potential strategy for modulating iNKT cell activity. By directly engaging iNKT TCRs, αGalCer/CD1d complexes can expand regulatory iNKT subsets and modulate broader T and B cell networks, suggesting a plausible role in ameliorating Dox-induced cardiotoxicity.

In this study, we investigated the frequency and distribution of iNKT subsets in the heart under homeostatic conditions and evaluated the therapeutic potential of αGalCer/CD1d complexes in Dox-induced cardiomyopathy. Our findings reveal that the heart harbors distinct iNKT subsets dominated by iNKT1 cells. Unexpectedly, TCR-targeted treatment with αGalCer/CD1d complexes provided only modest clinical improvements in a mouse model of Dox-induced cardiotoxicity, contrasting with prior reports. These results highlight the complexity of iNKT cell-mediated immune regulation in cardiac pathologies and emphasize the need for further exploration of targeted iNKT therapies for skewing to regulatory responses to prevent inflammation-induced cardiac damage.

## 2 Materials and Methods

### 2.1 Vertebrate animals

C57BL/6J male mice (10-weeks-old) were procured from the Jackson Laboratories (Strain #:000664, Bar Harbor, ME, USA) and were acclimated to the animal vivarium for 2 weeks. The mice were housed in cages with food and water *ad libitum* and maintained under a 12-h light/12- h dark photocycle in a temperature-controlled room. All animal experiments and procedures were approved by the University of Alabama at Birmingham Institutional Animal Care and Use Committee (UAB-IACUC) and the University Committee on Laboratory Animals (UCLA) at Dalhousie University.

### 2.2 Tetramers and antibodies

Mouse CD1d-KRN tetramers were procured from the NIH Tetramer Core Facility and used at a 1:1600 dilution. Fluorochrome-conjugated antibodies against mouse antigens were sourced from BD Biosciences (Mississauga, Canada) or Invitrogen (Carlsbad, CA), including CD3ε (500A2/145-2C11), CD4 (RM4.5), CD45 (30-F11), T-bet (4B10), PLZF (R17-908), GATA3 (L50-823), c-Maf (T54-853), and NK1.1 (PK136). Antibodies for surface staining were applied at a 1:200 dilution, while those for intracellular staining were used at a 1:100 dilution.

### 2.3 Heart tissue digestion and CD45+ immune cells purification

C57BL/6J mice were treated with either empty CD1d (healthy controls) or αGalCer/CD1d complexes once weekly for four weeks. Three days post-final injection, the hearts were harvested from each mouse as previously described (30) to quantify and characterize iNKT cells. The heart tissue was minced in 50 µL cold phosphate-buffered saline (PBS) in a petri dish and transferred to microcentrifuge tubes containing 1 mL of Dulbecco’s Modified Eagle Medium (DMEM) supplemented with 450 U/mL collagenase I (Roche Diagnostic, Indianapolis, IN), 60 U/mL hyaluronidase (Stemcell Technologies, Vancouver, Canada), and 60 U/mL DNase-I (Roche Diagnostic). The samples were incubated at 37°C for 45 minutes on a rocking shaker at 50 rpm. Following digestion, samples were vortexed for 20 seconds, placed on ice, and passed through a 40 µm cell strainer. The resulting cell suspensions were washed twice with cold Hank’s Balanced Salt Solution (HBSS) containing 0.2% BSA. Red blood cells were lysed using an Ammonium-Chloride-Potassium (ACK) lysis buffer. The remaining cells were washed and then subjected to CD45+ cell enrichment using the CD45+ Enrichment Kit (Stemcell Technologies), following the manufacturer’s instructions.

### 2.4 Flow cytometry

Single-cell suspensions obtained from CD45+ cell enrichment were stained with fixable viability dye eFluor 780 (Invitrogen) for 30 minutes in PBS at 4°C, followed by washing with FACS buffer (2% FBS and 20 mM sodium azide in PBS). To reduce non-specific binding, cells were incubated with Fc Block (CD16/CD32) for 15 minutes, followed by washing in FACS buffer. Surface staining was performed by incubating cells with fluorochrome-conjugated antibodies for 30 minutes at 4°C, after which they were washed in FACS buffer. Intracellular staining was carried out using the eBioscience FoxP3/Transcription Factor Staining Buffer Set, following the manufacturer’s protocol. Finally, cells were washed twice, fixed with 1% paraformaldehyde, and resuspended in FACS buffer. Flow cytometry data were acquired using a BD FACS Symphony cytometer and analyzed with FlowJo™ software version 10.8.1 (BD Biosciences).

### 2.5 CD1d complex production and loading of αGalCer into CD1d complexes

Mouse CD1d complexes were purified from CHO-S cell supernatants transduced with lentiviruses encoding β2m and CD1d. These CD1d complexes were engineered to include a 6×His and streptags, allowing for purification via affinity chromatography. To load αGalCer onto the mCD1d-empty complexes, KRN 7000 (Cayman Chemical, Ann Arbor, MI), dissolved in DMSO, was added to the mCD1d suspension at a molar ratio of 12:1 in the presence of PBS. The mixture was incubated for 3 hours at 37 °C, followed by an overnight incubation at 4 °C. The αGalCer- loaded CD1d monomers were then dialyzed in PBS for 12 hours and stored at 4 °C. The mCD1d concentration was quantified using the Bradford assay (Thermo Scientific).

### 2.6 Doxorubicin-induced cardiotoxicity and administration of αGalCer/CD1d complexes

Baseline echocardiography and body weights of experimental mice were analyzed. To induce chronic doxorubicin-mediated cardiotoxicity, mice were intraperitoneally administered with Doxorubicin hydrochloride (Dox) (3mg/kg body weight dissolved in saline; Cat# 2252, Tocris Bioscience) every other day for 2 weeks (total of eight Dox doses) (31). The healthy control mice were intraperitoneally administered with the same volume of saline. Since female C57BL/6J mice are reported to show resistance to doxorubicin-induced cardiac dysfunction (32), we only used male mice for the current study. The mortality in mice after Dox administration is shown in Supplementary Table 1.

After 4^th^ dose of Dox, Dox-treated mice were randomly divided and intraperitoneally administered with either CD1d-empty monomer or αGalCer/CD1d (20 µg/mouse/week, dissolved in PBS) for 7 weeks. The experimental animals were categorized in three groups: (1) Healthy control, (2) Dox+empty/CD1d, and (3) Dox+αGalCer/CD1d. The number of mice for each experiment is specified in the figure legends.

### 2.7 Echocardiography

Mice were anesthetized with a mixture of 1.5% isoflurane and oxygen (1 L/min) and transthoracic two-dimensional M-mode echocardiography was performed using Vevo2100 (VisualSonics, Canada) as described previously (33). Heart rate of the mice were maintained between 450-500 BPM. Left ventricular (LV) internal dimensions, LV anterior and posterior wall thicknesses, ejection fraction (EF), fractional shortening (FS), cardiac output, stroke volume, and other echocardiographic parameters were calculated using Vevo Lab software (version 5.8.0).

### 2.8 Body weight and organ weight measurements

Individual body weight of mice was recorded weekly after Dox treatment. At the endpoint, animals were euthanized, and the heart and spleen were removed immediately. The heart and spleen were washed in ice-cold PBS and weighed. The tibia length of experimental mice was also measured.

### 2.9 Histological analysis

At endpoint, animals were euthanized, and the heart tissues were removed immediately. Heart tissues were fixed in 10% neutral-buffered formalin and embedded in paraffin. Heart sections of 5 µm thickness were prepared and hematoxylin and eosin staining were done at the UAB Pathology Core. To evaluate cardiac fibrosis, Masson’s trichrome (Cat# HT15, Sigma-Aldrich) staining was performed as per the manufacturers’ instructions. The images of the LV region were captured under a Nikon Eclipse E200 microscope with NIS-Elements software version 4.60. A minimum of 200-250 cardiomyocytes from each mouse per group was traced for the measurement of cardiomyocyte area. For fibrosis measurement, 6-8 images of the LV region of Masson’s trichrome-stained sections were captured. The quantification of LV fibrosis and cross-sectional area of cardiomyocytes were determined using ImageJ (NIH) from captured images.

### 2.10 RNA extraction and quantitative real-time polymerase chain reaction (qRT-PCR) analysis

Total RNA from heart tissue was extracted using an RNA extraction kit (RNeasy Mini Kit, Cat# 74104, Qiagen) according to the manufacturer’s instructions. The RNA was reverse transcribed using the RevertAid First Strand cDNA Synthesis Kit (Cat# K1622, Thermo Fisher Scientific). Quantitative real-time PCR was performed in a QuantStudio 3 system (Applied Biosystems) using the PowerUp SYBR Green Master Mix (Cat# A25778, Thermo Fisher Scientific) according to the manufacturer’s instructions. Relative mRNA expression was normalized to GAPDH. Gene expression was determined by the comparative CT method (2^-ΔΔCT^) and was presented as fold change. The primer sequences of the genes are given in Supplemental Table.

### 2.11 Western blotting

To prepare whole cell lysate, the left ventricular tissues were lysed in RIPA buffer with a protease inhibitor cocktail. Protein concentrations were determined by Bradford assay (Bio-Rad, USA), and equal amounts of proteins were denatured in 4× Laemmli buffer. The denatured samples were resolved on denaturing SDS-PAGE gel (4-20%) and transferred to the PVDF membrane. The membranes were blocked with 4% bovine serum albumin (BSA) in tris-buffered saline with Tween-20 (TBS-T). The membranes were incubated with primary antibodies Bax (CST, Cat# 2272, 1:1000) and β-Tubulin (Proteintech, Cat# 66240-1-Ig, 1:10000), followed by HRP- conjugated secondary antibodies and visualized by an enhanced chemiluminescence (Pierce) detection system. Images of the blots were acquired using ChemiDoc™ Touch Imaging System (Bio-Rad, USA), and densitometric analyses were performed using ImageJ (NIH) software.

### 2.12 Terminal deoxynucleotidyl transferase-mediated dUTP nick-end labeling (TUNEL) assay

Cardiac cell death was determined using the In Situ Cell Death Detection Kit, Fluorescein (Cat# 11684795910, Millipore Sigma), according to the manufacturer’s instructions. Briefly, paraffin- embedded heart sections (5 μM thick) were deparaffinized and rehydrated through a graded series of ethanol and double-distilled water. The sections were treated with Proteinase K solution for 30 minutes at room temperature. After washing with PBS, the sections were incubated with a TUNEL reaction mixture for one hour at 37°C in a humidified chamber in the dark. The sections were washed with PBS and were mounted with VECTASHIELD^®^ PLUS antifade mounting medium with DAPI (Cat# H-2000, Vector Laboratories). The images were captured using an Olympus FV3000 confocal microscope at 40× magnification.

### 2.13 Statistical analysis

Data are presented using the mean±standard error of the mean (SEM). Normality of data was analyzed by Shapiro-Wilk test. Based on data distribution, One-way analysis of variance (ANOVA) with post-hoc Tukey or Kruskal-Wallis with post-hoc Dunn’s was performed to determine the statistical significance among multiple groups. Two-way repeated measures ANOVA for longitudinal data with post-hoc Tukey test for multiple pairwise comparisons was used for comparisons among multiple groups. Two-tailed Mann-Whitney U test was performed to determine the statistical significance between the two groups. A p-value of <0.05, indicated by * or # in the graphs, was considered statistically significant. Statistical analyses were performed using GraphPad Prism software version 10.2.0 (GraphPad Software Inc).

## 3 Results

### 3.1 αGalCer/CD1d treatment selectively expands cardiac iNKT1 cells

Building on our previous findings (20) demonstrating that iNKT-specific TCR-directed treatment expands liver-specific regulatory iNKT populations and ameliorates liver autoimmunity, we sought to determine whether similar effects extend to cardiac iNKT cells upon treatment with αGalCer/CD1d complexes. To this end, we treated C57BL/6J mice with 20 µg of αGalCer/CD1d or empty/CD1d complexes and assessed iNKT cell expansion in the heart three days post-treatment using flow cytometry. Our analysis revealed a striking three-fold expansion of total cardiac iNKT cells in mice treated with αGalCer/CD1d compared to those treated with unloaded CD1d complexes (Figure 1A). This result highlights the potential of TCR-directed therapeutics to significantly enhance iNKT cell populations within the heart.

**FIGURE 1.**
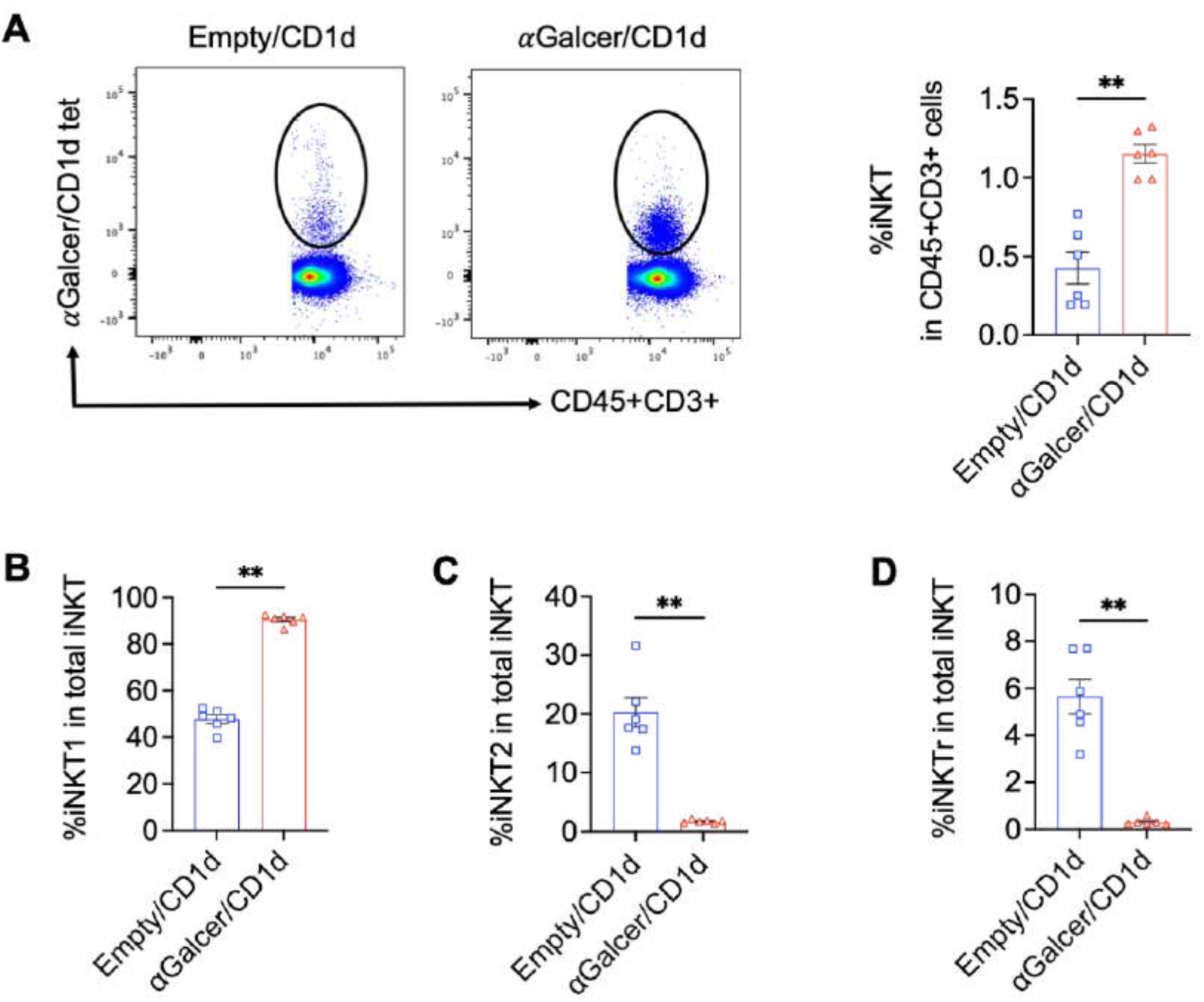
αGalCer/CD1d treatment preferentially expands cardiac iNKT1 cells. (A) Flow cytometric analysis of cardiac iNKT cell subsets in C57BL/6J mice injected with αGalCer/CD1d or Empty/CD1d complexes three days post-treatment. Representative dot plots (left panel) and bar graph (right panel) showing a significant expansion of total cardiac iNKT cells in αGalCer/CD1d-treated mice compared to the empty/CD1d. (B-D) Subset-specific analysis of cardiac iNKT cells revealed a preferential expansion of the pro-inflammatory iNKT1 subset (B), with a simultaneous reduction in iNKT2 (C) and iNKTr (D) subsets following αGalCer/CD1d treatment. Data are presented as mean±SEM (n=6/group) and were analyzed by the Mann-Whitney test. \**P*<0.05, \*\**P*<0.01 vs. Empty/CD1d control.

To further investigate the subset-specific effects of αGalCer/CD1d treatment under homeostatic conditions, we analyzed the distribution of iNKT subsets in the hearts of treated mice, using transcription factor and surface marker profiling. Interestingly, αGalCer/CD1d treatment selectively and significantly promoted the expansion of the pro-inflammatory iNKT1 subset in the heart compared to controls (Figure 1B). This finding was in contrast to our observations in liver tissues, where αGalCer/CD1d treatment supported a more balanced expansion of both regulatory iNKTR1 and iNKT2 subsets. Unexpectedly, αGalCer/CD1d treatment led to a marked reduction in cardiac iNKT2 and iNKTR subsets (Figure 1C, D), both of which are known to counteract the pro-inflammatory functions of iNKT1 cells. These observations suggest that soluble αGalCer/CD1d treatment skews the cardiac iNKT cell population towards a predominance of the iNKT1 subset, potentially altering the inflammatory balance within the heart. Flow cytometry gating strategy for different iNKT subsets cells is provided in Supplementary Figure S1.

### 3.2 Effect of αGalCer/CD1d administration on body and organ weights of experimental mice

We monitored the body weights of the control, Dox+empty/CD1d monomer-, and Dox+αGalCer/CD1d-administered mice weekly. The healthy mice continued gaining weight over time whereas Dox-administered mice progressively lost body weights as compared to the control mice (Figure 2A). At endpoint, the body weight of the Dox+empty/CD1d-administered mice were significantly lower than the control mice, and administration of αGalCer/CD1d did not reverse the loss of body weight (20.8±0.94g Dox+empty/CD1d, *P*<0.01; 21.0±0.52 g Dox+αGalCer/CD1d, *P*<0.01 vs. 31.3±1.12 g Control, Figure 2B).

**FIGURE 2.**
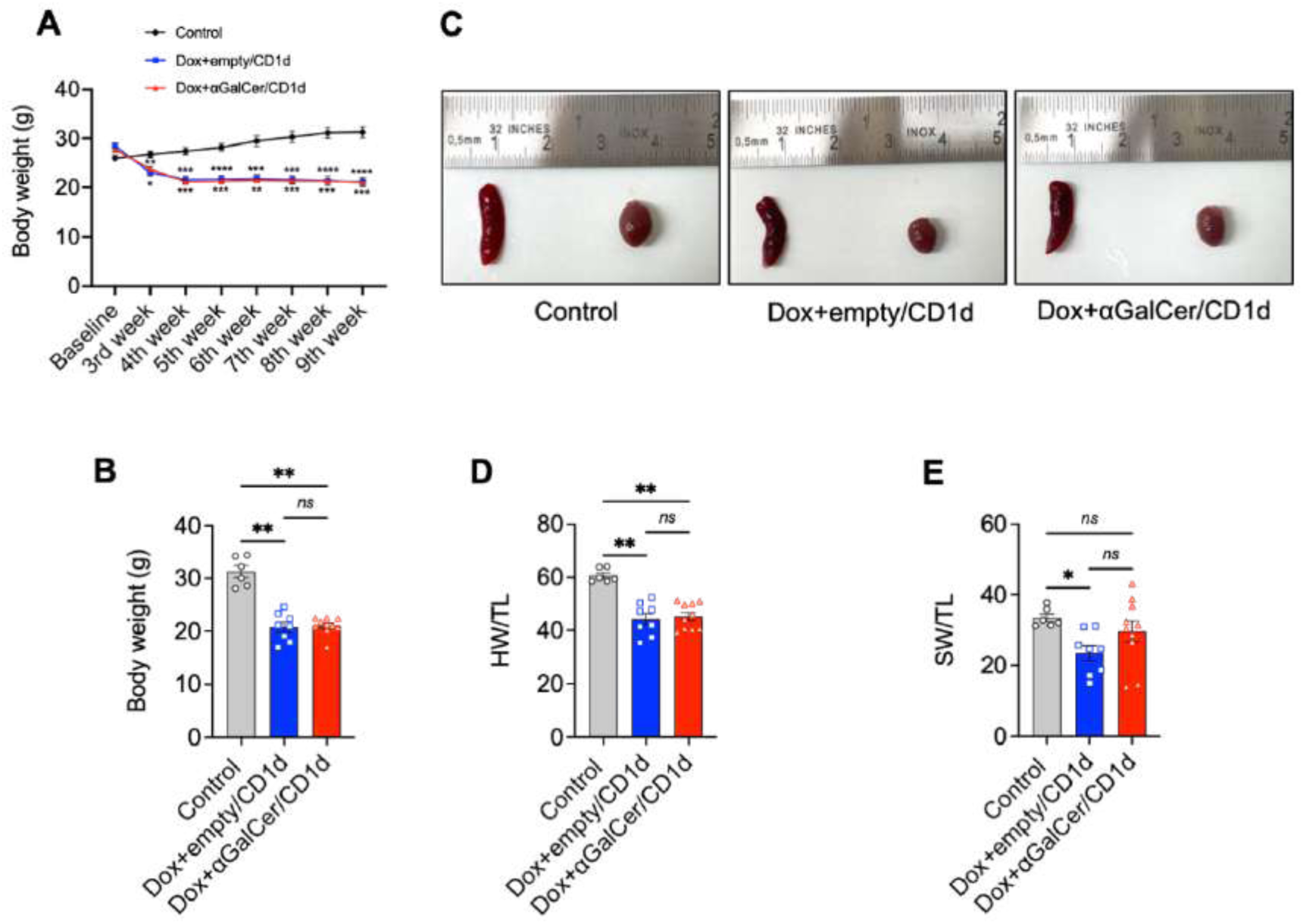
Effect of doxorubicin and αGalCer/CD1d administration on physiological parameters in experimental animals. (A) Weekly body weight measurement of the Control, Dox+empty/CD1d, and Dox+αGalCer/CD1d mice. (B) Body weight measurement at endpoint. (C) Representative images of the heart and spleen from the Control, Dox+empty/CD1d, and Dox+αGalCer/CD1d mice. (D) Heart weight to tibia length ratio. (E) Spleen to tibia length ratio. Data are presented as mean±SEM (n=6 for Control, n=8 for Dox+empty/CD1d, and n=10 for Dox+αGalCer/CD1d). Data in panel (A) were analyzed by Two-way repeated measures ANOVA with post-hoc Tukey. Data in panels (B) and (D) were analyzed by Kruskal-Wallis with post-hoc Dunn’s test. Data in panel (E) were analyzed by One-way ANOVA with post-hoc Tukey. \**P*<0.05, \*\**P*<0.01, \*\*\**P*<0.001, \*\*\*\**P*<0.0001 vs. Control.

The heart weight (HW) and spleen weight (SW) were measured. The hearts of Dox-administered mice were significantly smaller in size as compared to the control group (Figure 2C). When normalized to tibia length (TL), a significant reduction in HW/TL was observed in Dox-treated mice as compared to the control, suggesting cardiac damage (*P*<0.01; Dox+empty/CD1d, Dox+αGalCer/CD1d vs. Control). No difference in HW/TL ratio was observed between the Dox+empty/CD1d and Dox+αGalCer/CD1d groups (Figure 2D). The SW/TL was also significantly lower in Dox+empty/CD1d group compared to the control mice (*P*<0.05; Dox+empty/CD1d vs. Control). Interestingly, SW/TL ratio was comparable between the control and Dox+αGalCer/CD1d mice (Figure 2E).

### 3.3 Effect of αGalCer/CD1d administration on doxorubicin-induced cardiotoxicity and LV function

Cardiomyopathy and heart dysfunction induced by acute and chronic administration of doxorubicin are well established in rodent models (31, 32, 34). According to previous report (31), we administered a chronic dose of Dox (3mg/kg body weight every other for 2 weeks) to develop cardiac dysfunction in male C57BL/6J mice.

Cardiac function was measured at baseline by M-mode echocardiography and mice were randomly divided to the experimental groups. Baseline echocardiographic parameters are shown in Supplementary Table 2. Endpoint echocardiogram did not show any changes in ejection fraction, fractional shortening, and LV internal diameter at systole (LVID;s) after Dox administration (Figure 3A-C). The mice treated with Dox+empty/CD1d showed a marked decrease in LV internal diameter at diastole (LVID;d, *P*<0.05 vs. Control), LV mass (*P*<0.05 vs. Control), cardiac output (*P*<0.05 vs. Control), stroke volume (*P*<0.05 vs. Control), and end diastolic volume (EDV, *P*<0.05 vs. Control) as compared to the control mice (Figure 3D-H), confirming doxorubicin-induced cardiotoxicity. These changes were not reversed or restored by Dox+αGalCer/CD1d administration. No apparent changes were observed in end systolic volume (ESV) and anterior and posterior wall thicknesses among all treatment groups (Table 1).

**FIGURE 3.**
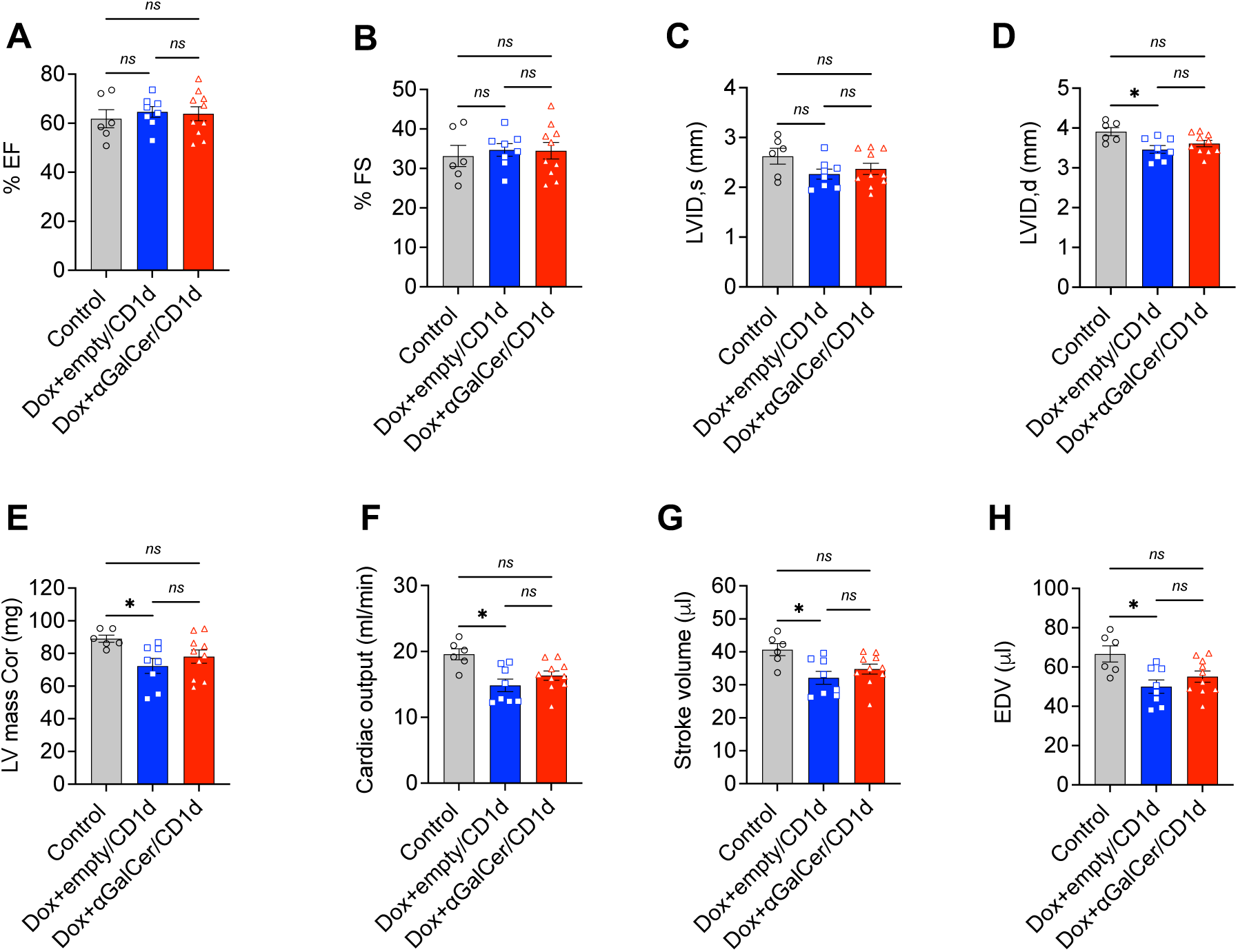
Effect of doxorubicin and αGalCer/CD1d administration on cardiac function. Evaluation of cardiac parameters by echocardiography. (A) Ejection fraction (%EF). (B) Fractional shortening (%FS). (C) Left ventricular internal diameter end systole (LVID;s). (D) Left ventricular internal diameter end diastole (LVID;d). (E) Left ventricular mass corrected (LV mass cor). (F) Cardiac output. (G) Stroke volume. (H) End diastolic volume (EDV). Data are presented as mean±SEM (n=6 for Control, n=8 for Dox+empty/CD1d, and n=10 for Dox+αGalCer/CD1d). Data in panels (A), (B), (C), (D), (E), (G), and (H) were analyzed by One-way ANOVA with post-hoc Tukey. Data in panel (F) were analyzed by Kruskal-Wallis with post-hoc Dunn’s test. \**P*<0.05 vs. Control.

**Table 1.** Echocardiographic parameters of Control, Dox+empty/CD1d, and Dox+αGalCer/CD1d-administered mice.

| Parameter | Control | Dox+empty/CD1d | Dox+αGalCer/CD1d |
| --- | --- | --- | --- |
| Heart rate (BPM) | 482.01±1.76 | 462.20±5.22 | 470.01±3.72 |
| ESV ( μl) | 25.98±3.64 | 17.91±2.06, P=0.13 | 20.29±2.40, P=0.31 |
| LVAW;s (mm) | 1.33±0.07 | 1.30±0.07, P=0.89 | 1.29±0.03, P=0.82 |
| LVAW;d (mm) | 0.86±0.04 | 0.90±0.04, P=0.81 | 0.89±0.02, P=0.93 |
| LVPW;s (mm) | 0.98±0.05 | 0.91±0.05, P=0.60 | 0.97±0.05, P=0.96 |
| LVPW;d (mm) | 0.71±0.02 | 0.66±0.04, P=0.59 | 0.69±0.04, P>0.99 |

Data are presented as mean ± SEM (n=6 for Control, n=8 for Dox+empty/CD1d, and n=10 for Dox+ αGalCer/CD1d). All parameters except LVPW;d were analyzed by One-way ANOVA with post-hoc Tukey. Data for LVPW;d were analyzed by Kruskal-Wallis with post-hoc Dunn's test. BPM: Beats per minute; ESV: End systolic volume; LVAW;s: left ventricular anterior wall end systole; LVAW;d: left ventricular anterior wall end diastole; LVPW;s: left ventricular posterior wall end systole; LVPW;d: left ventricular posterior wall end diastole. *P*-values are shown vs. Control group.

### 3.4 Evaluation of cardiac hypertrophic response in Dox-treated mice

We next investigated whether αGalCer/CD1d have any effect on cardiac hypertrophic response in Dox-administered mice. Both Dox+empty/CD1d and Dox+αGalCer/CD1d mice exhibited marked elevation in the expression of expression of atrial natriuretic peptide (*Nppa*, Figure 4A), B-type natriuretic peptide (*Nppb*, Figure 4B), and myosin heavy chain 7 (*Myh7*, Figure 4C) as opposed to the control group, confirming the onset of hypertrophic response. However, the levels of these markers were comparable between the Dox+empty/CD1d and Dox+αGalCer/CD1d groups. The expression of myosin heavy chain 6 (*Myh6*, Figure 4D) was similar among all the groups.

**FIGURE 4.**
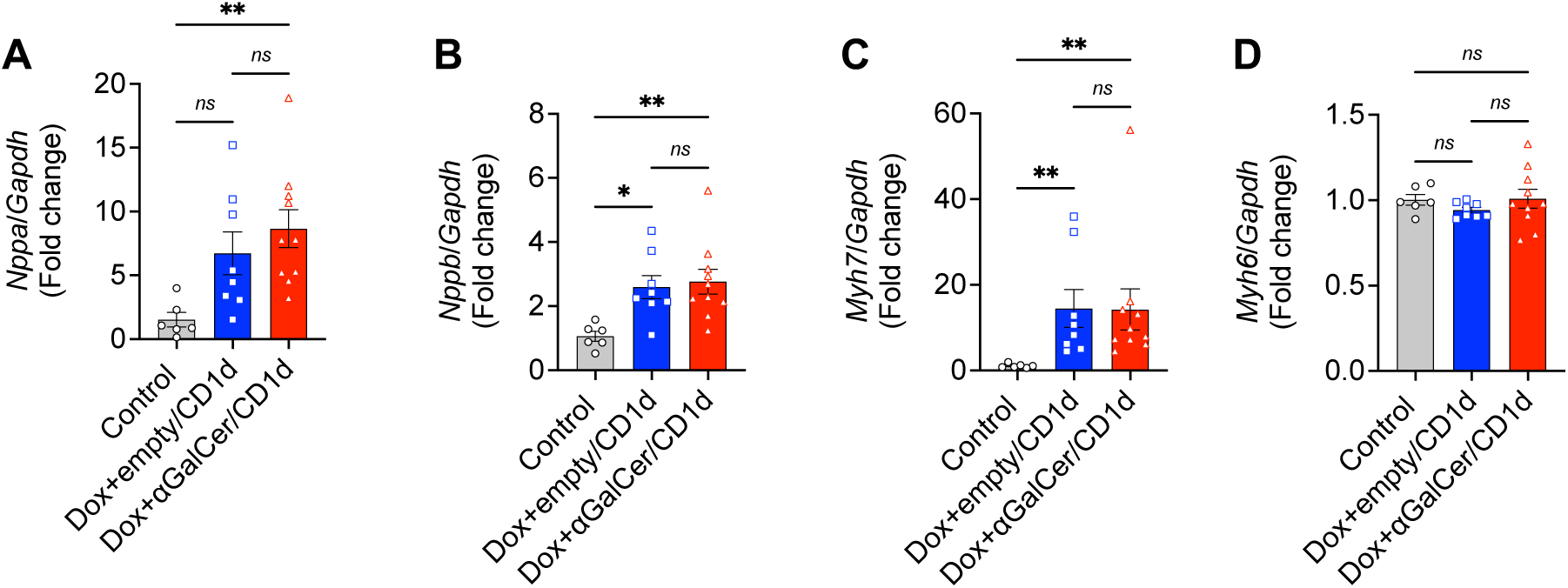
Effect of doxorubicin and αGalCer/CD1d administration on cardiac hypertrophic response. Cardiac mRNA expression of (A) atrial natriuretic peptide (*Nppa*), (B) B-type natriuretic peptide (*Nppb*), (C) myosin heavy chain 7 (*Myh7*), and (D) myosin heavy chain 6 (*Myh6*) normalized to *Gapdh*. Data are presented as mean±SEM (n=6 for Control, n=8 for Dox+empty/CD1d, and n=10 for Dox+αGalCer/CD1d). Data in panels (A), (B), and (D) were analyzed by One-way ANOVA with post-hoc Tukey. Data in panel (C) were analyzed by Kruskal-Wallis with post-hoc Dunn’s test. \**P*<0.05, \*\**P*<0.01 vs. Control.

We performed histological evaluation of cardiac structural changes. Cardiomyocyte cross-sectional area analysis of LV region did not exhibit any changes in the average cardiomyocytes size among the groups (Supplementary Figure S2).

### 3.5 Expression of fibrogenesis-related molecules in doxorubicin-treated mice after αGalCer/CD1d administration

Evaluation of fibrogenic markers in Dox+empty/CD1d and Dox+αGalCer/CD1d mice exhibited interesting expression pattern. The expression of *Col1a1*, *Col3a1*, and *Tgfb1* (TGF-β1) was reduced in the Dox+empty/CD1d group (Figure 5A-C) as compared to the control group. The expression of *Col1a1* was higher in the Dox+αGalCer/CD1d as compared to the Dox+empty/CD1d group, whereas the levels of *Col3a1* and *Tgfb1* were comparable between these two groups. Further, *Acta2* (α-SMA) and *Postn* (periostin) levels were comparable between the control and Dox+empty/CD1d groups but their expression was significantly increased in the Dox+αGalCer/CD1d mice (Figure 5D, E). The myocardial expression of *Ccn2* (connective tissue growth factor, CTGF) and *Tgfb2* (TGF-β2) was significantly higher in the Dox+empty/CD1d and Dox+αGalCer/CD1d mice compared to the control group and comparable between the Dox+empty/CD1d and Dox+αGalCer/CD1d groups (Figure 5F, G).

**FIGURE 5.**
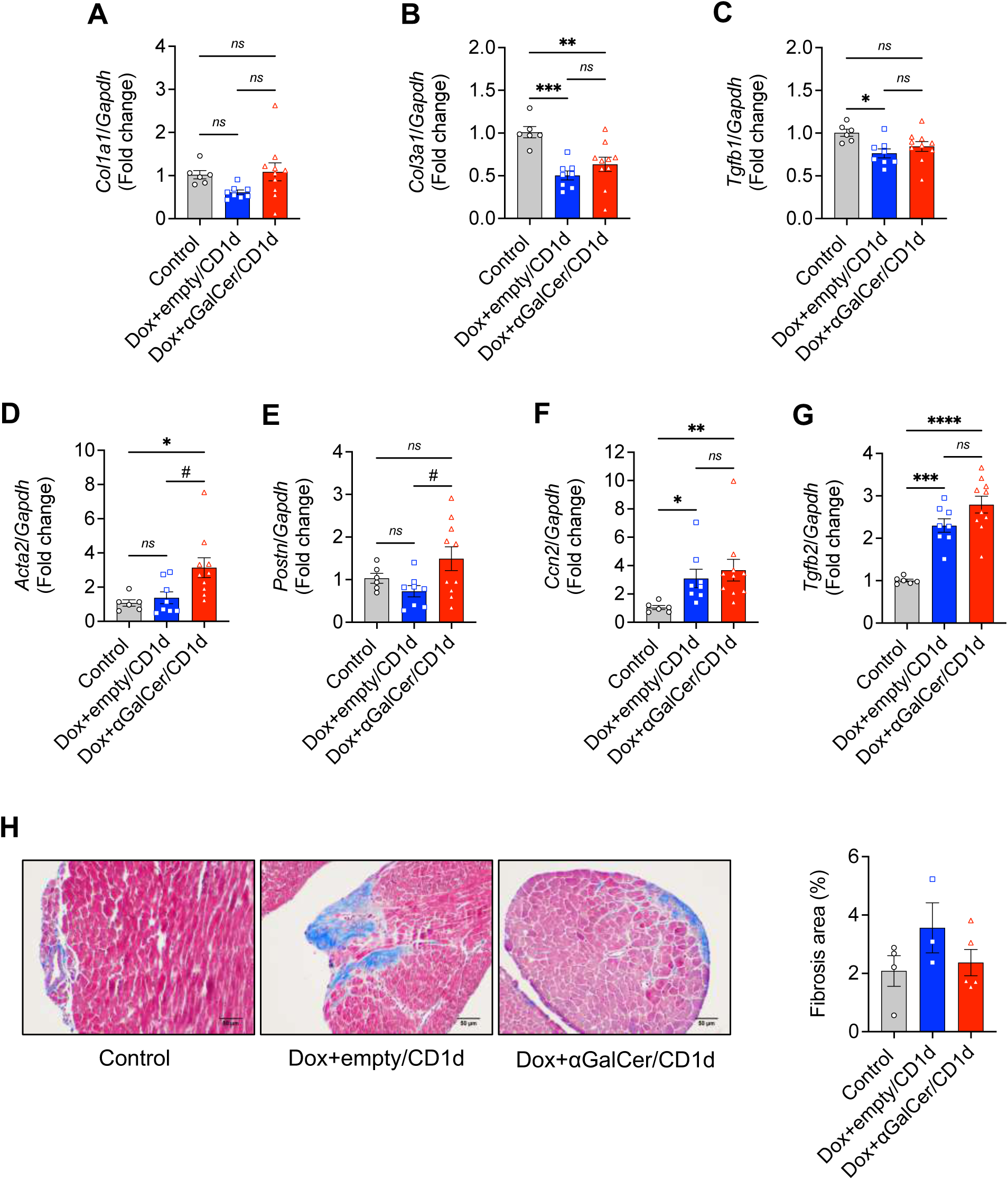
Expression of fibrogenic markers in doxorubicin-treated mice. Cardiac mRNA expression of (A) *Col1a1*, (B) *Col3a1*, (C) *Tgfb1*, (D) *Acta2*, (E) *Postn*, (F) *Ccn2*, and (G) *Tgfb2* normalized to *Gapdh*. Data are presented as mean±SEM (n=6 for Control, n=8 for Dox+empty/CD1d, and n=10 for Dox+αGalCer/CD1d). (H) Evaluation of collagen deposition in the hearts of doxorubicin-treated mice. Left panel: representative images of Masson’s trichrome staining in the heart tissues of the control, Dox+empty/CD1d, and Dox+αGalCer/CD1d groups. Right panel: quantification of % fibrotic area. Data are presented as mean±SEM (n=4 for Control, n=3 for Dox+empty/CD1d, and n=5 for Dox+αGalCer/CD1d). Data in panels (A), (B), (C), (E), (G) and (H) were analyzed by One-way ANOVA with post-hoc Tukey. Data in panels (D) and (F) were analyzed by Kruskal-Wallis with post-hoc Dunn’s test. \**P*<0.05, \*\**P*<0.01, \*\*\**P*<0.001, \*\*\*\**P*<0.0001 vs. Control. ^#^*P*<0.01 vs. Dox+empty/CD1d.

We performed Masson’s trichrome staining to evaluate the extent of cardiac fibrosis. Although we observed an increased collagen load in the Dox+empty/CD1d group as opposed to the control and Dox+αGalCer/CD1d groups (Figure 5H), a blind evaluation by a trained pathologist revealed that collagen deposition was within the normal limits in the control, Dox+empty/CD1d, and Dox+αGalCer/CD1d groups. These observations suggest that pathogenic fibrosis might not be established at the endpoint used in this study.

### 3.6 Effect of doxorubicin and αGalCer/CD1d administration on inflammatory response

At the endpoint, we examined the levels of inflammatory cytokines *Il6*, *Il1b*, and *Tnf* (TNF-α) in the heart tissues of the control, Dox+empty/CD1d, and Dox+αGalCer/CD1d mice. We did not observe any difference in the expression of these cytokines among the groups (Figure 6A-C). However, an early time-point evaluation of inflammatory markers could provide a comprehensive insight into the effect of αGalCer/CD1d administration in doxorubicin treatment.

**FIGURE 6.**
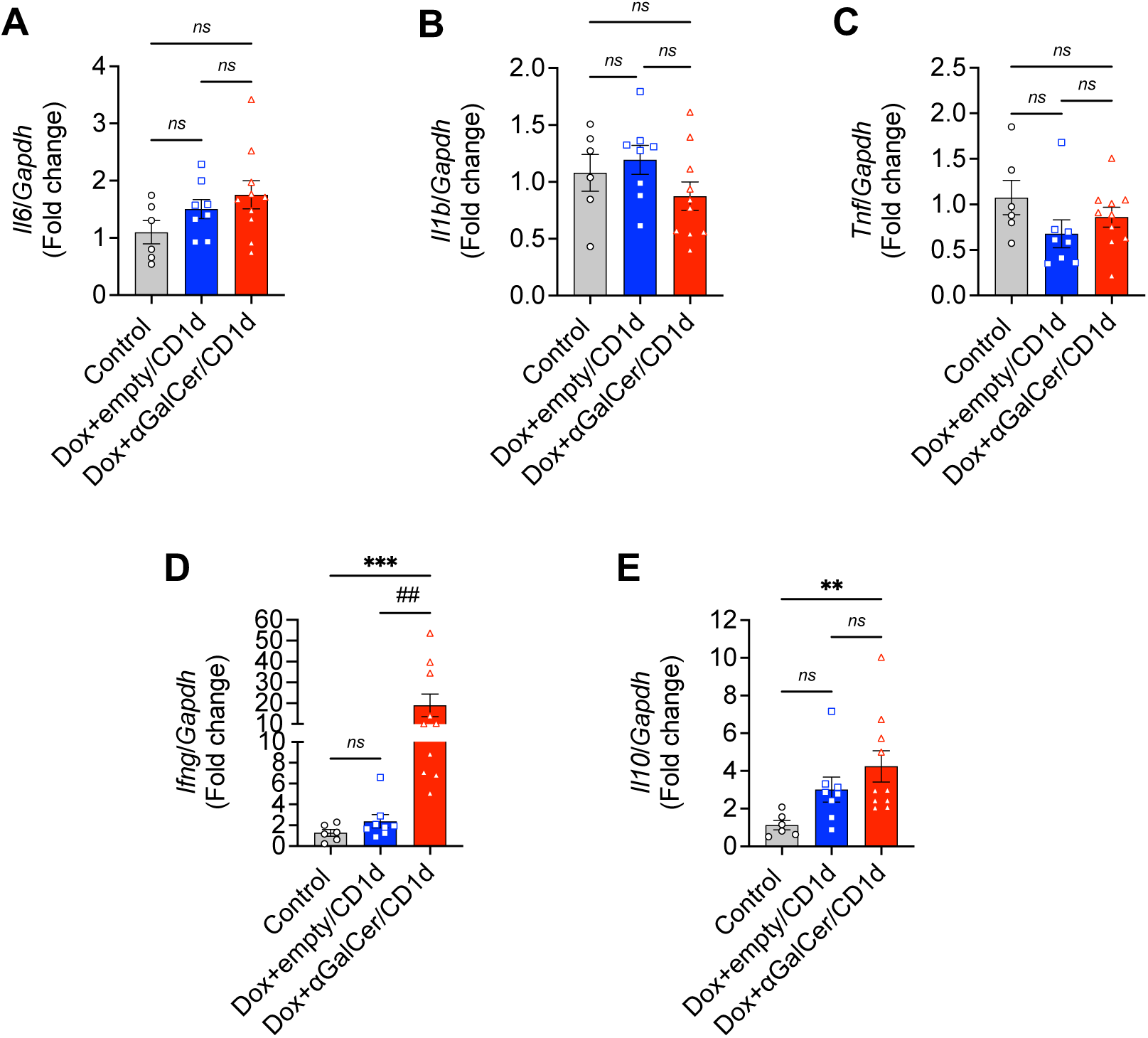
Expression of inflammatory cytokines in experimental groups. Myocardial gene expression of (A) *Il6*, (B) *Il1b*, (C) *Tnf* (TNF-α), (D) *Ifng* (IFN-γ), and (E) *Il10* normalized to *Gapdh*. Data are presented as mean±SEM (n=6 for Control, n=8 for Dox+empty/CD1d, and n=10 for Dox+αGalCer/CD1d). Data in panels (A) and (B) were analyzed by One-way ANOVA with post-hoc Tukey. Data in panels (C), (D), and (E) were analyzed by Kruskal-Wallis with post-hoc Dunn’s test. \*\**P*<0.01, \*\*\**P*<0.001 vs. Control. ^##^*P*<0.01 vs. Dox+empty/CD1d.

We next examined the cardiac expression of *Ifng* (IFN-γ) and *Il10* in the experimental mice. Interestingly, the expression of both *Ifng* and *Il10* was significantly upregulated in the Dox+αGalCer/CD1d group as compared to the control or Dox+empty/CD1d mice (Figure 6D-E), suggesting that these cytokines are predominant after αGalCer/CD1d administration. This observation also suggests the activation of iNKT cells in the heart tissue after αGalCer/CD1d treatment.

### 3.7 αGalCer/CD1d administration did not prevent doxorubicin-induced cardiac cell death

We next examined the effect of αGalCer/CD1d administration on Dox-induced myocardial cell death. The expression of pro-apoptotic molecule BAX was significantly upregulated in the Dox+empty/CD1d mice (Figure 7A). Similarly, a higher number of TUNEL-positive nuclei were observed in the left ventricles of the Dox+empty/CD1d mice than in the control group (Figure 7B). However, contrary to previously published reports (35), αGalCer/CD1d administration could not attenuate doxorubicin-induced cardiac cell death.

**FIGURE 7.**
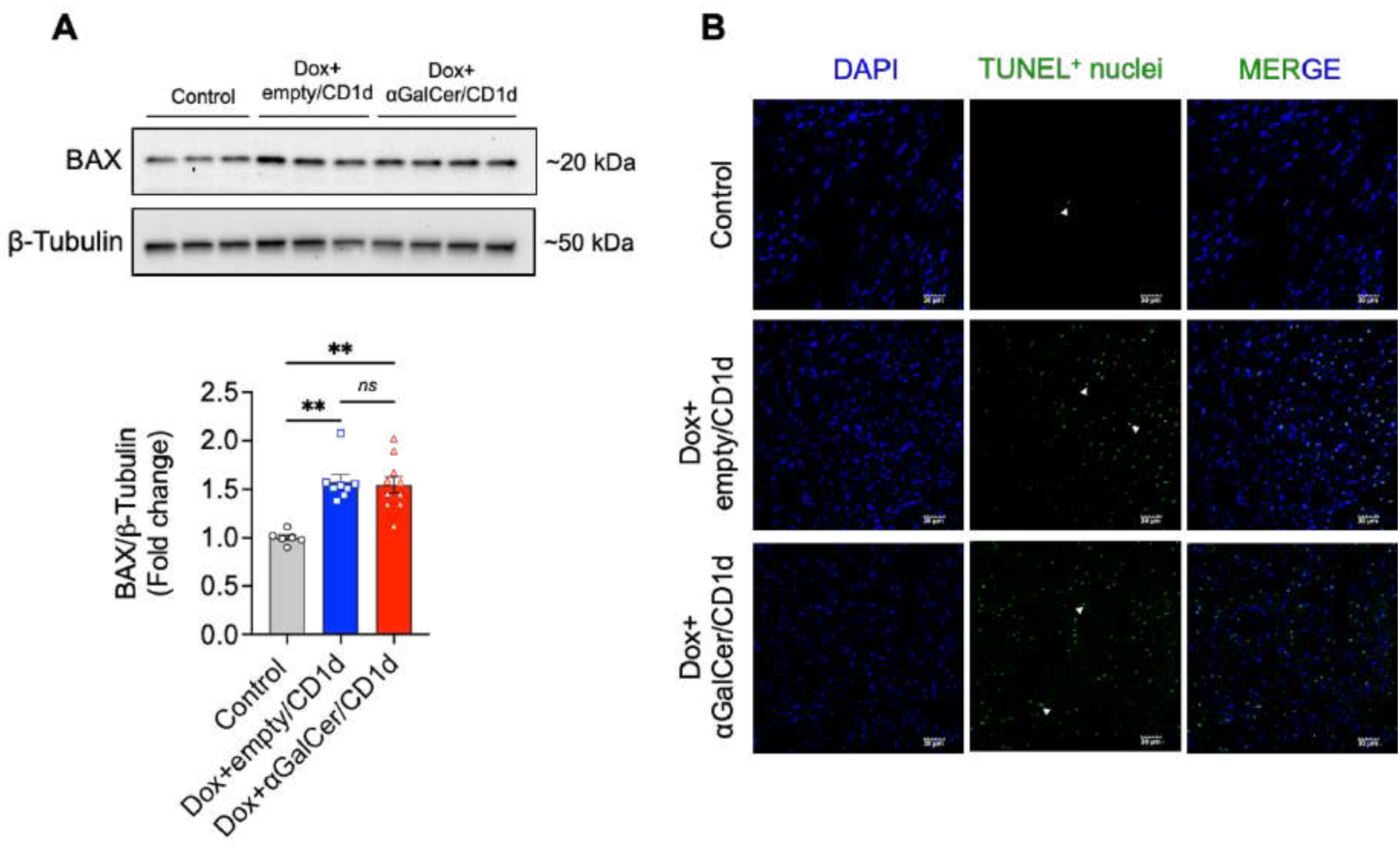
Effect of doxorubicin and αGalCer/CD1d administration on cardiac cell death. (A) Cardiac protein expression of BCL2-associated X (BAX). Upper panel: representative Western blot images of BAX and β-Tubulin; lower panel: densitometric analysis of BAX expression normalized to β-Tubulin. Data are presented as mean±SEM (n=6 for Control, n=8 for Dox+empty/CD1d, and n=10 for Dox+αGalCer/CD1d) and were analyzed by Kruskal-Wallis with post-hoc Dunn’s test. \*\**P*<0.01 vs. Control. (B) Representative images of TUNEL staining in the left ventricles of Control, Dox+empty/CD1d, and Dox+αGalCer/CD1d mice. White arrowheads indicate the TUNEL-positive nuclei in the cardiac sections.

## 4 Discussion

The role of tissue-specific iNKT subsets in various organs such as the liver, lungs, spleen, and lymph nodes is well-established (6, 14–22); however, their distribution and response to TCR-directed immunotherapies in the heart remain largely unexplored. Our study identifies that the healthy heart predominantly contains the iNKT1 subset, followed by iNKT2, and the recently discovered regulatory subset iNKTR1 (6, 20). These findings highlight the potential roles of these subsets in maintaining cardiac homeostasis and defense. The predominance of iNKT1 suggests the heart is primed for rapid inflammatory responses against infections, while iNKT2 and iNKT17 subsets likely contribute to anti-inflammatory functions and protection against bacterial and fungal pathogens, respectively (6). Additionally, the presence of iNKTR1 suggests a regulatory mechanism mitigating inflammation, which may be vital given the constant exposure of the heart to mechanical and physiological stressors.

TCR-directed immunotherapies have shown promise in modulating immune responses in autoimmune and inflammatory diseases (20, 36–40). Soluble peptide-MHC complexes, for example, induce tolerance and reduce inflammation in autoimmune diseases (28, 41–44). We hypothesized that similar approaches, such as treatment with soluble αGalCer/CD1d complexes, would favor the expansion of anti-inflammatory subsets like iNKTR1 and iNKT2 at the expense of the pro-inflammatory iNKT1 subset. Unexpectedly, our findings indicate that αGalCer/CD1d treatment selectively expanded iNKT1 at the cost of iNKT2 and iNKTR subsets. This skewing towards a pro-inflammatory response warrants further investigation, particularly in the context of inflammatory cardiac diseases such as Dox-induced cardiotoxicity. Future studies will aim to delineate the dynamics of iNKT subset alterations during Dox-induced cardiotoxicity and their response to TCR-directed therapeutics.

Previous studies have demonstrated the cardioprotective effects of αGalCer in conditions such as myocardial infarction and ischemia/reperfusion injury (45). These benefits were attributed to reduced T cell infiltration, decreased pro-inflammatory cytokines (e.g., IL-1β, TNF-α), and increased anti-inflammatory mediators such as IL-10 and IL-4 (46). Furthermore, αGalCer (αGC) treatment has been shown to mitigate Dox-induced left ventricular dysfunction and fibrosis by promoting M2 macrophage polarization (47). Interestingly, these studies used a single high dose (20 mg/kg) of Dox, where cardiac manifestations are more severe. Since the high-dose model of doxorubicin causes severe cardiac atrophy, we used a lose-dose (3 mg/kg) model of Dox-induced cardiotoxicity, which is more clinically relevant. We observed a significant decrease in cardiac output, stroke volume, LV mass, and other parameters confirming the Dox-inflicted cardiotoxicity. However, we did not observe any changes in the ejection fraction and fractional shortening, suggesting that the cardiac changes are less severe than a high-dose Dox. To evaluate whether αGalCer/CD1d treatment could prevent the systolic dysfunction similar to αGC administration, we intraperitoneally administered αGalCer/CD1d in Dox-treated mice for 7 weeks. Surprisingly, in our study, αGalCer/CD1d treatment did not provide similar cardioprotective effects; however, it did not worsen the condition. A recent study by Sada et al. has shown that simultaneous administration of αGC with Dox did not attenuate cardiotoxicity, whereas pre-treatment of αGC could protect against Dox-induced cardiotoxicity and cell death (35). These observations suggest that even though αGalCer/CD1d complexes directly engage with iNKT TCRs, bypassing lipid antigen presentation by local APCs, pre-treatment of these complexes could be necessary for their cardioprotective effects. These findings emphasize the importance of understanding the interplay between iNKT subsets, antigen presentation, and the cardiac immune microenvironment in shaping therapeutic outcomes. Future studies with pre- and continuous administration of αGalCer/CD1d could provide more insights into its mechanism in Dox-induced cardiomyopathy.

Previous studies have shown a cardioprotective effect of iNKT cells in different disease models. Takahashi et al. reported that cardiac function and remodeling worsen in iNKT cell-deficient *Jα18* knockout mice after pressure-overload cardiac injury (48). Cardiac fibrosis and myocardial hypertrophy were exacerbated in the iNKT cell-deficient *Jα18* knockout mice after transverse aortic constriction (TAC) (48). However, conflicting results have been reported regarding the antifibrotic effect of iNKT cells in Dox-induced cardiomyopathy (35, 47). Obata et al. showed a protective effect of αGC administration on cardiac fibrosis in Dox-induced cardiotoxicity (47). On the contrary, Sada et al. did not observe any reduction in myocardial fibrosis after αGC administration and speculated that myocardial fibrosis may not be present in their model (35). In the current study, the pathologist could not identify pathological fibrosis at the endpoint, although there was a trend toward increased fibrosis in the hearts of empty/CD1d-treated mice compared to αGalCer/CD1d-treated and untreated healthy mice. Nonetheless, further studies with extended durations and larger sample sizes are warranted to conclusively determine the antifibrotic role of iNKT cells in the heart.

Significantly, we observed increased expression of *Ifng* mRNA in the damaged heart tissue of Dox-treated mice, correlating with the expansion of the pro-inflammatory iNKT1 subset in αGalCer/CD1d-treated C57BL/6J mice. Despite this, there was no exacerbation of cardiac damage, likely due to the concurrent upregulation of *Il10* mRNA. Interestingly, both iNKT2 and iNKTR subsets, known producers of IL-10, were reduced following KRN/CD1d treatment. This suggests the potential involvement of an unidentified regulatory iNKT subset contributing to IL-10 secretion and mitigating inflammatory damage. Identifying and characterizing this subset will be pivotal for understanding the regulatory mechanisms at play and optimizing therapeutic strategies.

Variations in TCR signaling across different tissues and immune contexts may account for the observed variability in iNKT cell responses. For instance, αGalCer has demonstrated efficacy in treating autoimmune diseases like type 1 diabetes, systemic lupus erythematosus, and multiple sclerosis (11, 12, 49–52) but has been less effective in liver autoimmunity (20). In our previous work, αGalCer/CD1d-coated nanoparticles successfully expanded liver-specific regulatory iNKT subsets, termed Maf^+^LiNKTR1, which mediated tolerance and suppressed liver inflammation (20).

These regulatory cells mediate their suppressive effects by inducing a robust regulatory network involving B cells and APCs. These peptide-MHC-coated nanoparticles provide sustained TCR signalling and additional tolerogenic cues for the differentiation of regulatory T cells (37). In contrast, soluble αGalCer/CD1d complexes may provide suboptimal signals, leading to incomplete differentiation of regulatory subsets in the heart, as observed with free αGalCer treatment in liver autoimmunity (20). Future studies should explore whether nanoparticle-based delivery systems could more effectively expand cardiac-specific regulatory iNKT subsets to control inflammatory conditions like Dox-induced myocarditis.

In conclusion, our findings reveal that the heart predominantly harbors the pro-inflammatory iNKT1 subset, with smaller populations of anti-inflammatory iNKT2, iNKT17, and iNKTR1 subsets. While αGalCer/CD1d treatment favored iNKT1-derived *Ifng* expression, it did not exacerbate cardiac damage, likely due to compensatory IL-10 production by an unidentified regulatory subset. These results underscore the complexity of iNKT subset interactions within the cardiac immune microenvironment and highlight the potential of targeting iNKT cells for therapeutic benefit. Moving forward, the development of modified TCR-directed immunotherapeutics holds significant promise. Strategies such as nanoparticle-based αGalCer/CD1d delivery systems or tolerogenic cell-based therapies could provide a more precise and context-specific approach to modulating iNKT responses. These advanced therapeutics could be leveraged to enhance immune defense mechanisms in conditions like sepsis and infections or to promote immune regulation in inflammatory cardiac diseases, including Dox-induced myocarditis, myocardial infarction, and autoimmune myocarditis.

## Data availability statement

The data that support the findings of this study are available in the article and supplementary information.

## Author contributions

All authors contributed substantially to the study. CSU and PK conceptualized the initial project and provided intellectual insights and critical appraisal. SS and RHN performed experiments with contributions from HBK, DKS, and AP. SS, RHN, PK, and CSU analyzed the data. SS, RHN, CSU, and PK interpreted the data. SS, RHN, and CSU wrote and revised the manuscript draft with input and assistance from PK, HBK, DKS, and AP.

## Acknowledgements

This work was supported by the Canadian Institutes of Health Research (CIHR) through a Tier 2 Canada Research Chair award (2021-00215, Dr. Umeshappa), Dalhousie University start-ups fund (Dr. Umeshappa), National Institutes of Health (NIH) (HL138023, Dr. Krishnamurthy), and US Department of Defense grant (PR220330, Dr. Krishnamurthy). Dr. Singh was supported by the American Heart Association postdoctoral fellowship (916497). Dr. Nanjundappa received support from the Dalhousie Medical Research Foundation (DMRF) and the American Association of Immunologists (AAI) Fellowship. Mr. Kolla was supported by the DMRF I3V Graduate Studentship. Dr. Chauhan received funding through the I3V Hubel and IWK Postdoctoral Fellowships (award 1028897).

