## Supplementary files for "αGalCer/CD1d treatment induces pro-inflammatory iNKT1-associated immune responses without aggravating doxorubicin-induced cardiotoxicity"

### Supplementary Data

**Supplementary Table S1. Mice mortality in the control, Dox+empty/CD1d, and Dox+αGalCer/CD1d mice**

| Group | Starting mice number | Death | Death rate (%) |
|---|---|---|---|
| Control | 6 | 0 | 0 |
| Dox+empty/CD1d | 10 | 1 | 10 |
| Dox+αGalCer/CD1d | 10 | 0 | 0 |

**Supplementary Table S2. Baseline echocardiographic parameters of experimental animals**

| **Parameter** | **Baseline** |
|---|---|
| Heart rate (BPM) | 471.66±6.47 |
| Ejection fraction (%) | 66.98±1.76 |
| Fractional shortening (%) | 36.72±1.38 |
| LVID;s (mm) | 2.39±0.07 |
| LVID;d (mm) | 3.77±0.05 |
| LV mass corrected (mg) | 89.96±3.44 |
| Cardiac output (ml/min) | 19.12±0.57 |
| Stroke volume (µl) | 40.69±1.37 |
| ESV (µl) | 20.29±1.58 |
| EDV (µl) | 60.98±2.08 |
| LVAW;s (mm) | 1.35±0.05 |
| LVAW;d (mm) | 0.93±0.03 |
| LVPW;s (mm) | 1.02±0.03 |
| LVPW;d (mm) | 0.73±0.02 |

Data are presented as mean±SEM (n=12). BPM: Beats per minute; LV: left ventricle; LVID;s: left ventricular internal diameter end systole; LVID;d: left ventricular internal diameter end diastole; ESV: End systolic volume; EDV: End diastolic volume; LVAW;s: left ventricular anterior wall end systole; LVAW;d: left ventricular anterior wall end diastole; LVPW;s: left ventricular posterior wall end systole; LVPW;d: left ventricular posterior wall end diastole.

**Supplementary Table S3: Mouse primer sequences used for real-time quantitative PCR**

| **Mouse primers** | | |
|---|---|---|
| **Gene** | **Forward primer (5'-3')** | **Reverse primer (5'-3')** |
| *Nppa* | GCTTCCAGGCCATATTGGAG | GGGGGCATGACCTCATCTT |
| *Nppb* | GAGGTCACTCCTATCCTCTGG | GCCATTTCCTCCGACTTTTCT |
| *Myh6* | GCCCAGTACCTCCGAAAGTC | GCCTTAACATACTCCTCCTTGTC |
| *Myh7* | ACTGTCAACACTAAGAGGGTCA | TTGGATGATTTGATCTTCCAGGG |
| *Col1a1* | GCTCCTCTTAGGGGCCACT | CCACGTCTCACCATTGGGG |
| *Col3a1* | CTGTAACATGGAAACTGGGGAAA | CCATAGCTGAACTGAAAACCACC |
| *Tgfb1* | CTCCCGTGGCTTCTAGTGC | GCCTTAGTTTGGACAGGATCTG |
| *Acta2* | CCCAGACATCAGGGAGTAATGG | TCTATCGGATACTTCAGCGTCA |
| *Postn* | CCTGCCCTTATATGCTCTGCT | AAACATGGTCAATAGGCATCACT |
| Ccn2 | GGGCCTCTTCTGCGATTTC | ATCCAGGCAAGTGCATTGGTA |
| *Tgfb2* | CTTCGACGTGACAGACGCT | GCAGGGGCAGTGTAAACTTATT |
| *Il6* | TAGTCCTTCCTACCCCAATTTCC | TTGGTCCTTAGCCACTCCTTC |
| *Il1b* | GCAACTGTTCCTGAACTCAACT | ATCTTTTGGGGTCCGTCAACT |
| *Tnf* | CCCTCACACTCAGATCATCTTCT | GCTACGACGTGGGCTACAG |
| *Ifng* | AGACAATCAGGCCATCAGCA | TGGACCTGTGGGTTGTTGAC |
| *Il10* | GCTCTTACTGACTGGCATGAG | CGCAGCTCTAGGAGCATGTG |
| *Gapdh* | AGGTCGGTGTGAACGGATTTG | TGTAGACCATGTAGTTGAGGTCA |

**Supplementary Fig. S1**

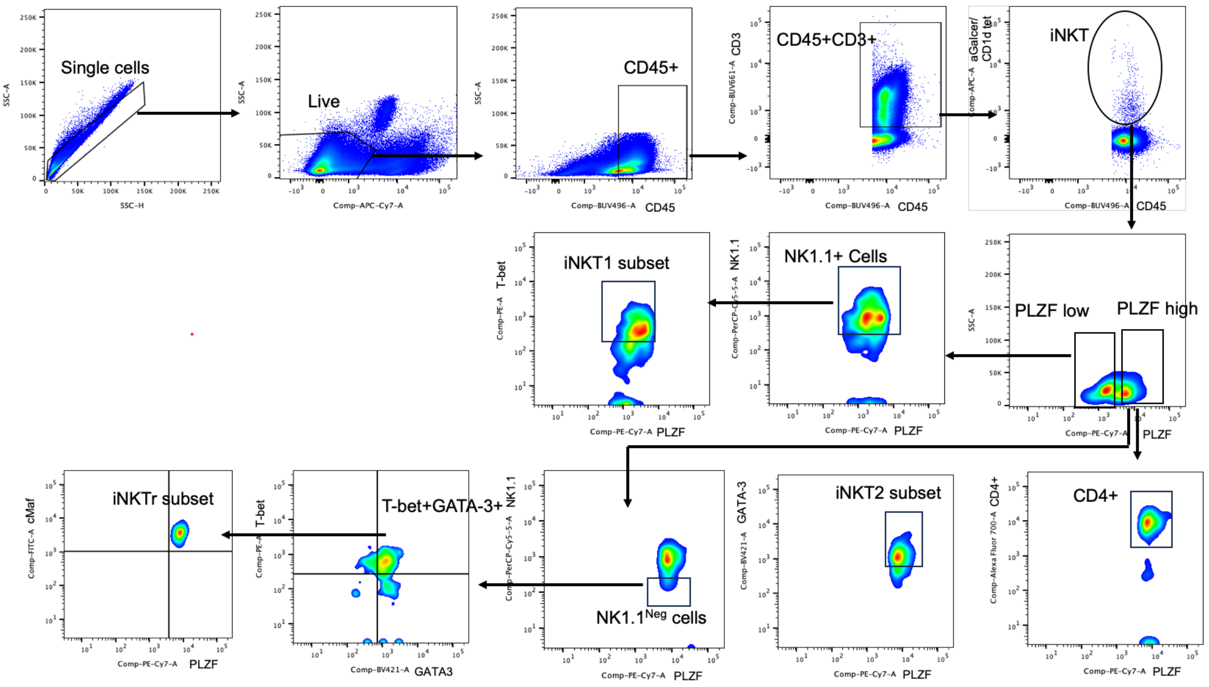


**Fig. S1.** Flow cytometry gating strategy for different iNKT subsets cells. iNKT cells were defined as CD45$^{+}$CD3$^{+}$ αGalcer/CD1d tetramer$^{+}$. iNKT1 subset was identified as PLZF$^{low}$NK1.1$^{+}$T-bet$^{+}$; iNKT2 subset was identified as PLZF$^{high}$CD4$^{+}$GATA-3$^{+}$; iNKTr subset was identified as PLZF$^{high}$NK1.1$^{neg}$T-bet$^{+}$ GATA-3$^{+}$ cMaf$^{+}$.

**Supplementary Fig. S2**

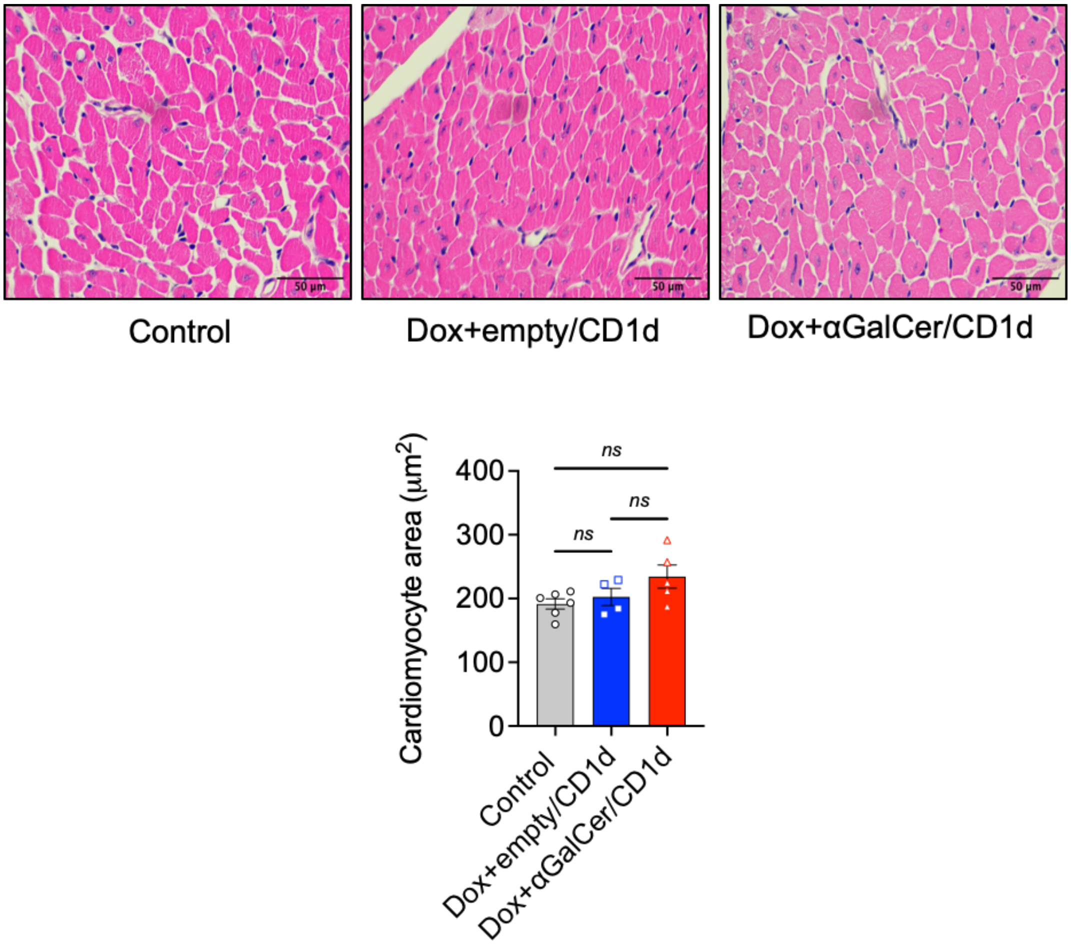


**Fig. S2.** Quantification of cardiomyocyte cross-sectional area in hematoxylin and eosin-stained sections. Upper panel: representative images of hematoxylin and eosin staining; Lower panel: quantification of cardiomyocyte area. Data are presented as mean±SEM (n=6 for Control, n=4 for Dox+empty/CD1d, and n=5 for Dox+αGalCer/CD1d). Data were analyzed by One-way ANOVA with post-hoc Tukey.

**Supplementary Fig. S3**

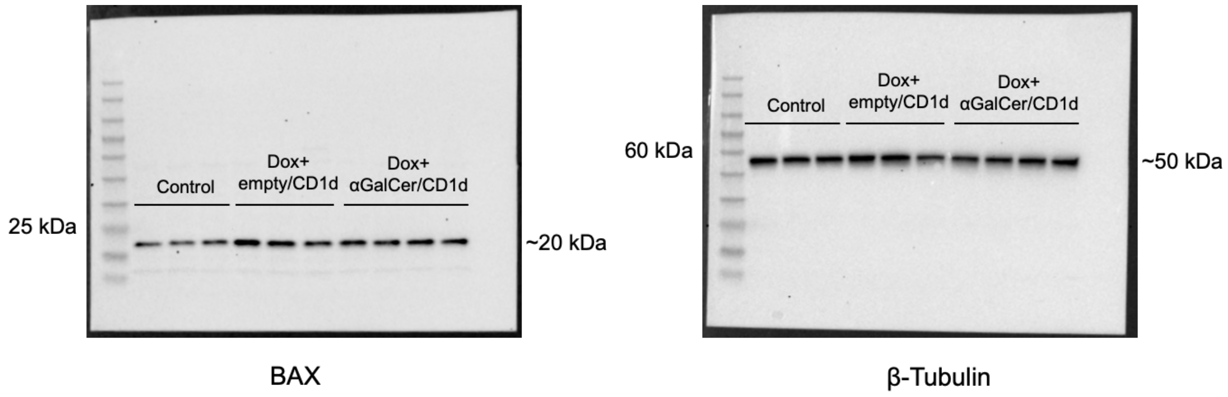


**Fig. S3.** Unedited blots for BAX and β-Tubulin for the representative Fig. 7A.
